# Community attitudes on genetic research of gender identity, sexual orientation, and mental health

**DOI:** 10.1101/685982

**Authors:** Taylor R. Thomas, Dabney Hofammann, Brooke G. McKenna, Anna I.R. van der Miesen, Mark A. Stokes, Peter Daniolos, Jacob J. Michaelson

## Abstract

Biological sex is an important factor in mental health, and a non-binary view of how variation in sex and gender influence mental health represents a new research frontier that may yield new insights. The recent acceleration of research into sexual orientation, gender identity, and mental health has generally been conducted without sufficient understanding of the opinions of sexual and gender minorities (SGM) toward this research. We surveyed 768 individuals, with an enrichment of LGBTQ+ stakeholders, for their opinions regarding genetic research of SGM and mental health. We found that the key predictors of attitudes toward genetic research specifically on SGM are 1) general attitudes toward genetic and mental health research 2) tolerance of SGM and associated behaviors 3) non-cisgender stakeholder status and 4) age of the respondent. Non-heterosexual stakeholder status was significantly associated with increased willingness to participate in genetic research if a biological basis for gender identity were discovered. We also found that non-stakeholders with a low tolerance for SGM indicated their SGM views would be positively updated if science showed a biological basis for their behaviors and identities. These findings represent an important first step in understanding and engaging the LGBTQ+ stakeholder community in the context of genetic research.

## 1 Introduction

Biological sex interacts with other risk factors in ways that are often strongly predictive of health outcomes. From immune response [15] to heart disease [16] and depression [14], sex is a key variable that contextualizes risk factors [27] and can yield biological insights into disease mechanisms. What is less understood is whether a continuous (rather than binary) view of sex and gender can deliver additional explanatory power in studies of human health and disease. Involving sexual and gender minorities (SGM, or more commonly referred to as the LGBTQ+ community –lesbian, gay, bisexual, transgender, and queer) can provide an important view of gender and sexuality that is not restricted to a strictly binary perspective when considering sex differences in human health. Of particular interest are neuropsychiatric conditions that have strong sex biases and have shown evidence of enrichment for non-heterosexuality and gender variance. For instance, autism is highly male-biased (4:1) [10] and is also enriched for gender dysphoria [24] [13], while anorexia nervosa is highly female-biased (8:1) [23] but is enriched for gay and bisexual men [9]. More generally, there has been extensive study into the higher prevalence of neuropsychiatric conditions in the LGBTQ+ community [22]. While it is possible that the increased prevalence of neuropsychiatric conditions in the LGBTQ+ community are due primarily to sociocultural factors like prejudice, stigma, discrimination, and rejection [17], few studies have investigated potential biological factors underlying these connections, especially working with the understanding that sex and gender are more complex, continuous factors.

As the tools for genomic research have become more accessible, fields beyond medicine, including social science, are increasingly appealing to genetic data in the search for explanatory factors of human behavior and identities. This trend reflects an entry into the “genomics to society” phase of a tripartite goal laid out in 2003 by the National Human Genome Research Institute [7]. With this transition comes an urgent need to understand the perspectives and concerns of both the general public and the groups being studied (what we refer to as stakeholder groups). This is particularly true in the case of sexual orientation, gender identity, and their potential connections to aspects of mental health, which have been subject to recent genetic investigations [18][26]. While genetics cannot fully explain these sensitive and often stigmatized [1] aspects of individual identity, a greater understanding of the genetic and biological contributions to these phenomena may reduce public stigma while also advancing scientific understanding of the complex relationships between sex, gender, and risk for neuropsychiatric conditions. This dual potential can only be achieved through partnership between scientists and the stakeholder community.

Recent technology and data availability has created more opportunities for expansive research into sexual orientation and gender identity, and many research groups are utilizing these advances. The results of these studies can have profound implications for millions of people (4.5% of the United States population belongs to the LGBTQ+ community [2]). For example, in 2018 a research group developed a deep neural network that used social media facial photographs to estimate sexual orientation with an accuracy of 81% in men and 74% in women [25]. This study was not governed by an institutional review board (IRB) and showed no evidence of engagement with stakeholders with respect to study design, implementation, or communication of the results. With the proliferation of social media and ease of access to facial photographs, the potential negative consequences of research in this area deserves thoughtful consideration and partnership with stakeholders. Likewise, although genetic research into human behavioral traits like sexual orientation has been proceeding for many years, it has recently accelerated, utilizing large and often publicly available data sets. Recently, researchers described the largest genome-wide associated study of sexual orientation to date by mining two massive genetic data sets, the UK Biobank and 23AndMe [12]. Genetic research into non-cisgender identities is gaining traction as well, with the International Gender Diversity Genomics Consortium being organized in 2018, their goal being to study the heritable, polygenic component of gender identity [19]. Other recent work examined variants in sex-hormone signaling and found significant differences in repeat-length variants and single nucleotide polymorphisms in transgender women compared to cisgender men [11]. This research has strong potential to deepen the understanding of the genetic component of complex human behaviors and experiences, but it also has potential to be misconstrued or abused. Historically there has been little consideration for LGBTQ+ voices and their opinions and concerns in this research area. Therefore, the objective of this study was twofold: first, to obtain a systematic, data-driven assessment of attitudes related to genetic research of sexuality and gender identities, and secondly to give a voice in the scientific literature to stakeholder groups and use their opinions to help inform the research that affects them.

## 2 Methods

This study was approved the University of Iowa’s Institutional Review Board (IRB #201611784). The survey was built on the Qualtrics platform. The majority of respondents were recruited via mass email to the University of Iowa, as well as recruitment through social media.

### 2.1 Online survey

The survey was designed to capture the respondent’s knowledge of and views on genetic research broadly, as well as genetic research into mental health, neuropsychiatric conditions, sexual orientation, and gender identity. We also asked questions regarding the respondent’s opinions on non-heterosexuality and non-cisgender identities. Opinion data was collected on a 5-point Likert scale. Considering our main goal was to understand the opinions of people in specific communities, we collected in-depth data regarding their own sexual orientation, gender expression, and gender identity. In addition, we collected basic demographic information. Demographic data for sexual orientation, gender identity, religion, race, and ethnicity were collected with the respondent able to select multiple values to describe themselves. A demographic summary is displayed in Table 2. Survey respondents were also asked to arrange continuously-adjustable sliders (representing femininity, masculinity, and “other”, which they could name themselves in a free text box) in a way that best described their gender identity. At the end of the survey, respondents were asked open-ended, free text questions in which they were able to detail their opinions and concerns, and were given the option to provide contact information to be contacted about future research opportunities. The complete survey is available in Supplemental File 1.

**Table 1:** Key definitions for terms used.

| Definitions |  |
| --- | --- |
| Transgender | an umbrella term that describes an individual who does not identify or exclusively identify with their recorded sex |
| Cisgender | term describing an individual who identifies with the gender that is consistent with their recorded sex |
| Recorded sex | sex recorded at time of birth based on physiological and anatomical sex characteristics; has also been referred to as natal sex or assigned sex |
| Gender | the behavioral norms for each sex that emerge at the population level, driven by a combination of biological, social, and cultural influences |
| Gender identity | the relation of an individual to gender norms that is most consistent with that individual's feelings, perspectives, and behaviors |
| Gender expression | the way in which an individual shows their gender identity through physical appearance, behavior, and interests |
| Sexual orientation | a component of an identity that includes a person's sexual and emotional attraction to another person, as well as the behavior and/or social affiliation that may result from this attraction |
| Non-heterosexuality | an encompassing sexual identity term for those who are not strictly heterosexual |
| Gender variance | the discrepancy between an individual's gender identity and the gender typical of their recorded sex |
| Sexual and gender minorities (SGM) | an encompassing term for people who are not heterosexual and/or not cisgender |
| Stakeholder | an individual who has an investment in a group and is affected by their involvement in the group |

**Table 2:**
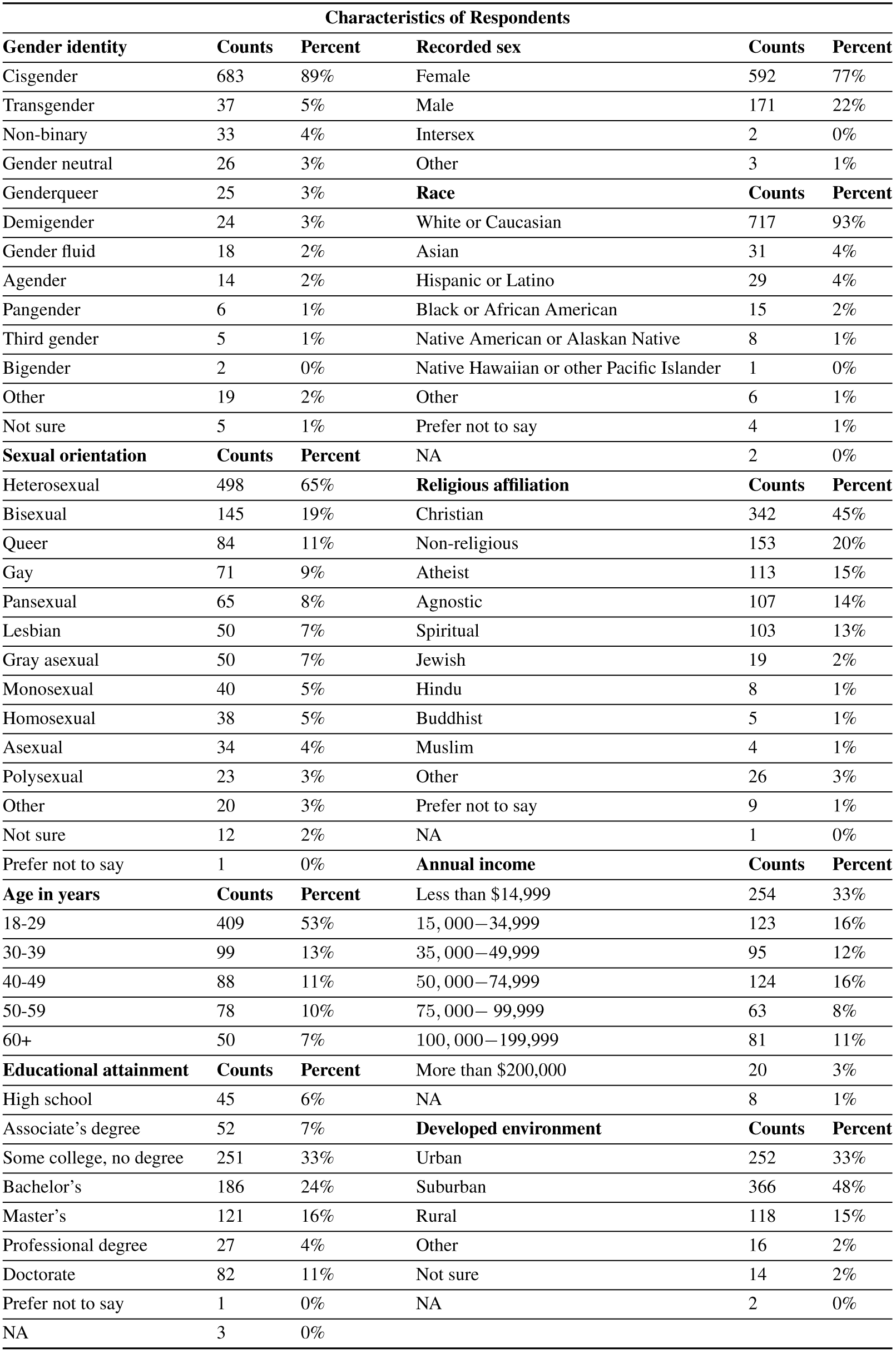
Counts and percentage of respondents who endorsed a demographic characteristic. For gender identity, sexual orientation, race, and religious affiliation, individuals were able to endorse multiple identities. For example, the respondent could have selected bisexual, pansexual, and queer, which contributes a count to each of those three categories.

### 2.2 Imputation of missing data

Tabular survey data was extracted from Qualtrics and analyzed in R. Surveys completed in less than two minutes or with excessive missing data were discarded. Overall, 1.2% of the data was missing, and missing data was imputed using a nearest-neighbors weighted mean approach as implemented in the `rfImpute()` function in the `randomForest` package for R, using a response variable defined by all possible combinations of recorded sex, gender identity (cisgender or non-cisgender), and sexual orientation (heterosexual or non-heterosexual). This has the effect of imputing missing data with heavier weights on individuals that share the same combinations of recorded sex and stakeholder status as the subject of the imputation. A total of *N=768* responses were used in the subsequent analyses.

### 2.3 Statistical analysis

The imputed tabular survey data was used to test the association of individual survey items, as well as a composite score (a linear combination of all these items weighted by 1 if it represented an optimistic statement and -1 if it represented a pessimistic statement) with the explanatory factors shown in Figure 1. The glm function in R was used to carry out these tests in a multiple linear regression framework, with `family=“quasipoisson”` for Likert-scale items, `family=“binomial”` for binary items, and `family=“gaussian”` for the composite score. The test statistics shown in Figure 1 are those from models where each row was modeled as a function of all columns included together. Correcting for multiple testing was performed using the Benjamini-Yekutieli procedure [6] for false discovery rate (FDR), which is valid under arbitrary assumptions, including correlated hypotheses. In addition, boxplots in Figure 1 show variance explained for each column variable across each row survey item or composite score.

**Figure 1:**
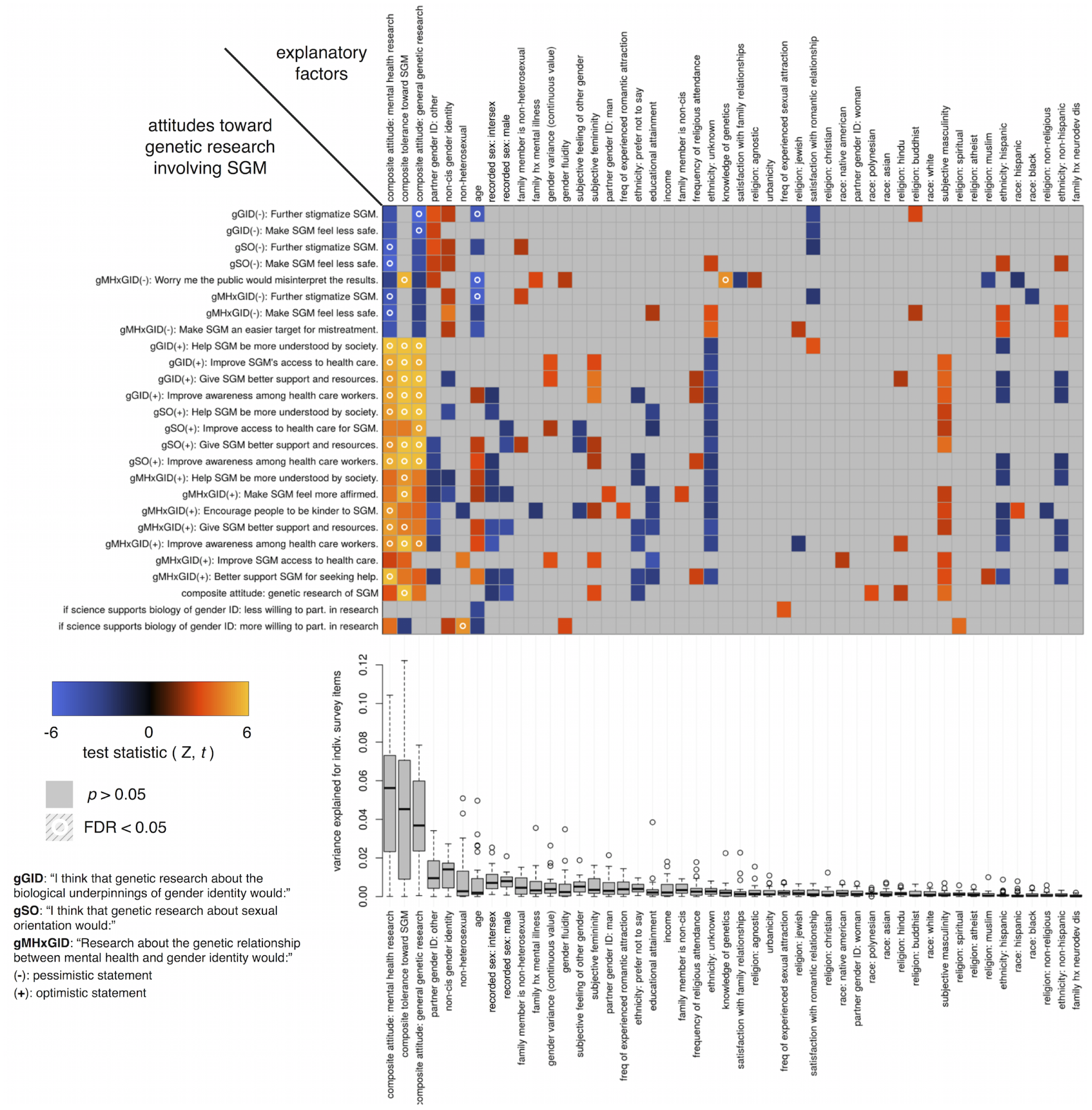
The statistical associations between survey items related to attitudes toward genetic research involving SGM (rows) and explanatory factors (columns). Explanatory factors (columns) are ordered by the mean variance explained (boxplots below) over the considered survey items. Negative associations are in blue, positive associations are in orange, and non-significant associations are in gray. Boxes with a circle represent associations that survive multiple testing correction.

### 2.4 Power

A key question is whether our survey sample is sufficiently powered to detect whether stakeholder status (i.e., non-cisgender identity or non-heterosexual orientation) is a significant contributor to opinions on genetic research involving SGM. We used the `pwr.r.test()` function from the `pwr` package for R to calculate power, which yielded estimates of 0.8-0.98 assuming an effect size (Pearson’s *r* for this test) of 0.1-0.14, suggested by the observed 1-2% variance explained by non-cisgender identity (see boxplots in Figure 1).

### 2.5 Gender space and continuous-valued gender variance

In addition to providing categorical descriptors of gender identity that respondents could choose to endorse, we provided three continuously-adjustable sliders (0-1 for each of femininity, masculinity, and “other”, which respondents could re-name if they wished), and asked respondents to arrange the sliders in a way they felt was most consistent with their gender identity. The values from these sliders comprise a three-dimensional gender space, where each respondent can be described by a triplet coordinate of [femininity, masculinity, other] (Fig. 2). We calculated a scalar-valued gender variance score by taking each respondent’s gender coordinates, and calculating the Euclidean distance to a gender datum of [1,0,0] if the respondent reported being a recorded female, and [0,1,0] if the respondent reported being a recorded male. For this particular analysis, those who reported “intersex” as their recorded sex (*N=2*) were excluded.

**Figure 2:**
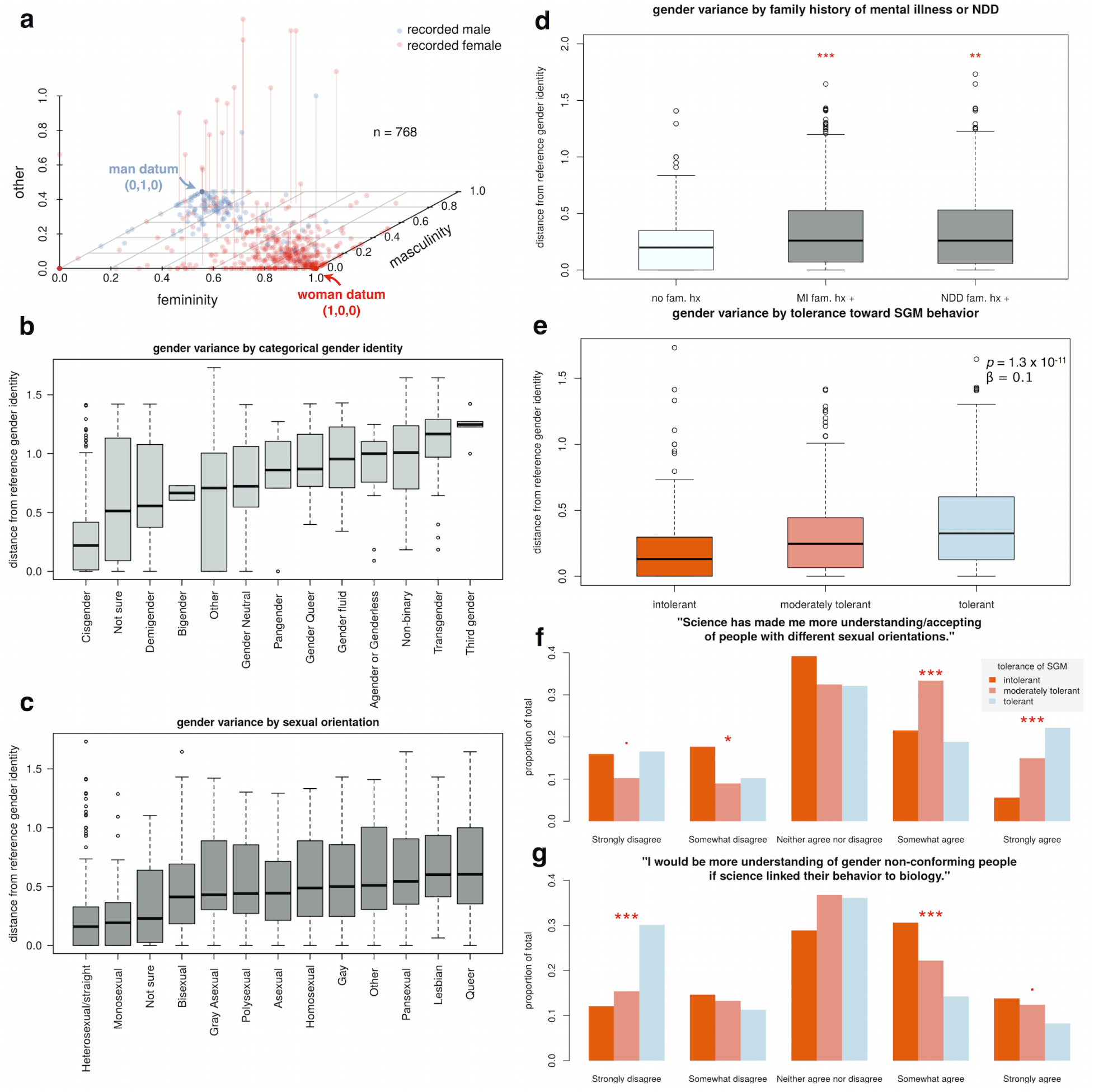
A three-dimensional gender identity space (a) was used to calculate a scalar-valued gender variance score, which is the Euclidean distance from the expected reference gender identity (woman or man datum) based on the recorded sex of the respondent. Scalar gender variance corresponded in a largely expected way to categorical descriptors of gender identity (b), and sexual orientation (c). We found that respondents with a family history of either mental illness or neurodevelopmental disorders (NDD) showed greater gender variance (d). Those who indicated greater tolerance of SGM also showed greater gender variance (e). When reflecting on previous shifts in personal acceptance of non-heterosexual orientations, respondents with higher current levels of tolerance attributed some of that shift to science (f). When considering future attitudes toward gender non-conformity, those who currently display the lowest levels of tolerance were significantly more likely to endorse science as a potential avenue toward increased personal acceptance (g). Significance key:. = p < 0.1; * = p < 0.05; ** = p < 0.01; *** = p < 0.001

### 2.6 Tolerance indicator

In order to evaluate how overall tolerance of non-heterosexuality and gender variance influenced opinions towards genetic research of sexual orientation and gender identity, we developed a tolerance indicator. This was built using Likert scale responses to the statements presented in Table 3. To facilitate a grouped analysis of tolerance as it relates to other measures, we binned respondents into three groups, which we labeled as “intolerant” (bottom quartile), “moderately tolerant” (interquartile range) and “tolerant” (top quartile).

**Table 3:**
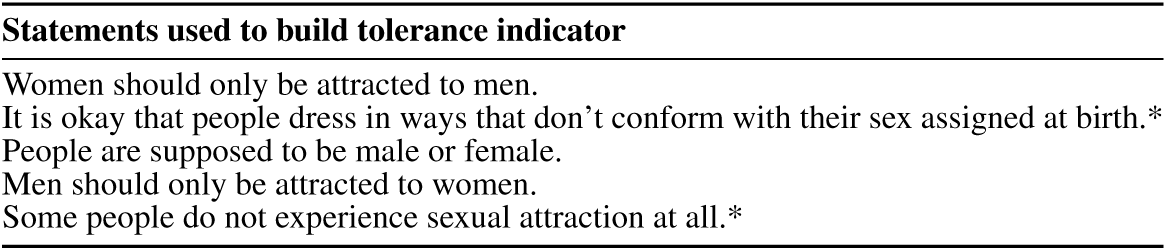
Respondents answered on a Likert scale to the following statements which were used to build a non-heterosexuality and gender variance tolerance indicator. Statements marked with an asterisk were scored positively towards higher tolerance.

## 3 Results

The sample characteristics of our respondents are described in Table 2. We identified non-cisgender individuals by a respondent who did not exclusively identify as cisgender and selected at least one of the other gender identities. We identified non-heterosexual individuals by a respondent who did not exclusively identify as heterosexual and selected at least one of the other sexual orientations. The sample shows a significant enrichment of the LGBTQ+ community: 11% (*N=85*) vs. 0.5% nationally (P < 0.001) reporting non-cisgender identity and 35% (*N=270*) vs. 4.1% nationally (P < 0.001) reporting non-heterosexual orientation. In addition, stakeholder status was significantly (P<0.001, Wilcoxan test) and positively associated with the volume of feedback in the free text fields (see Table 4 for representative examples).

**Table 4:**
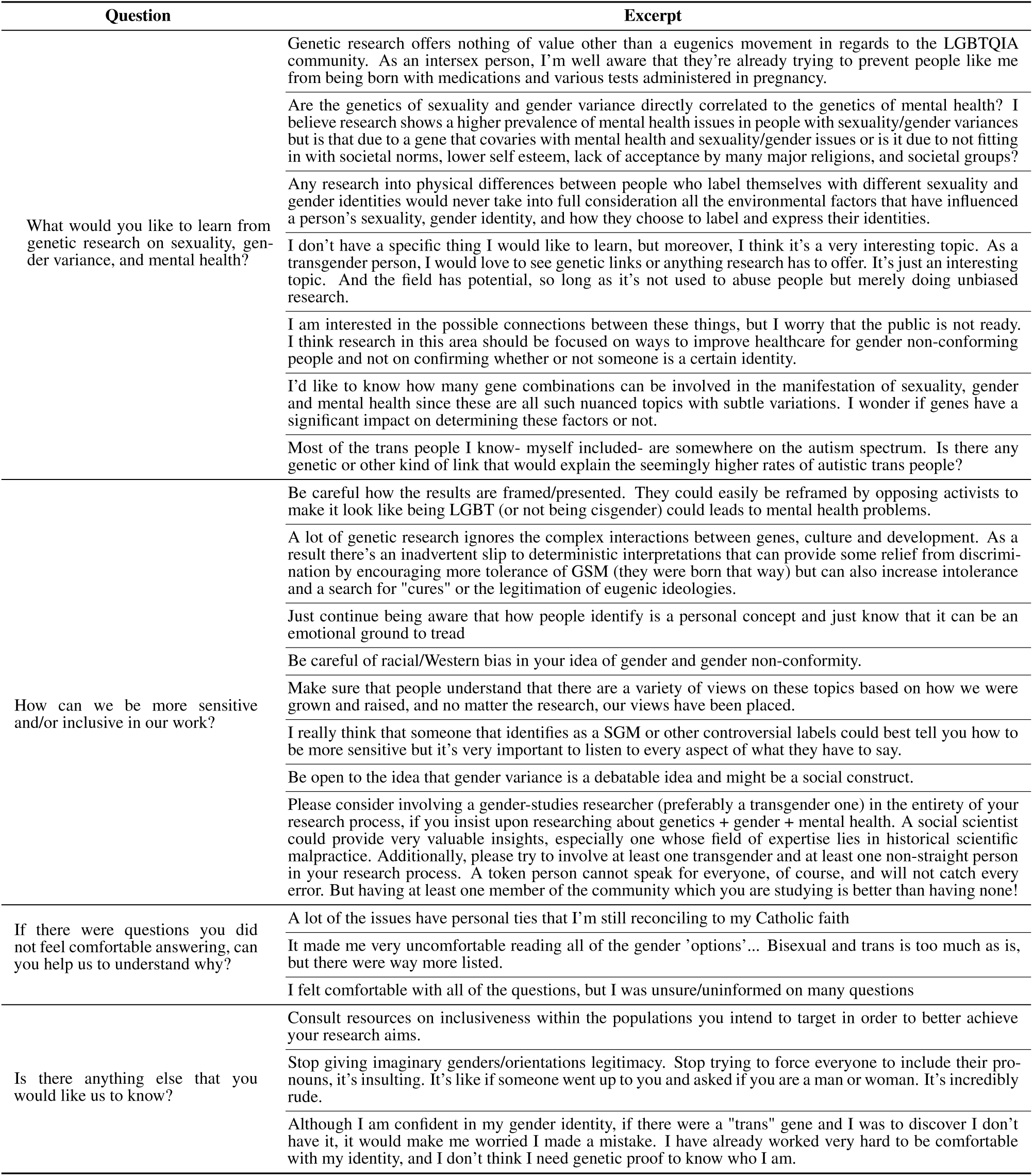
Representative excerpts taken from free text responses. Respondents were given the option to answer all, any, or none of the questions. We selected responses that were representative of a frequently expressed thought, or that we found to be particularly compelling.

### 3.1 Key drivers of attitudes toward genetic research on SGM

We found that attitudes toward genetic research specifically on SGM are most strongly predicted by broader attitudes toward mental health and genetic research in general (Fig. 1). Respondents who expressed reservations about mental health or genetic research in general were significantly more likely to express reservations or pessimism about genetic research on SGM. Similarly, respondents who endorsed the value of genetic or mental health research generally were also more likely to view genetic research on SGM positively.

We used the tolerance indicator (see Table 3) to build a composite non-heterosexuality and gender variance tolerance score. This composite tolerance score was significantly and positively associated with attitudes on genetic research involving SGM, meaning that those who were more accepting of SGM and associated behaviors were more likely to view genetic research involving SGM favorably. A notable exception to this pattern was observed on the item that expressed concern that the public would misinterpret genetic findings involving SGM, where tolerance was significantly (FDR < 0.05) associated with positive endorsement of the concern (Fig. 1). This item also showed significant associations with age (negative, FDR < 0.05), and objectively measured knowledge of genetics (positive, FDR < 0.05), suggesting that concern about public misinterpretation of the results of genetic research on SGM is expressed most strongly among younger respondents who are more tolerant and more conversant in genetic concepts.

To a lesser extent, stakeholder status related to non-cisgender identity (for the respondent themselves and/or for their romantic partner) trends toward concern about genetic research involving SGM. None of these associations survived correction for multiple hypothesis testing. Closer examination of these associations revealed that extreme heterogeneity of opinion within stakeholder groups, rather than small effect sizes, is the key factor preventing stronger associations of attitudes with stakeholder status. In any case, power analyses (see methods) suggest that our sample is sufficiently powered to detect stakeholder effects when they exist.

Despite traces of concern about genetic research on SGM, both non-cisgender and non-heterosexual respondents indicated that they would be more likely to participate in genetic research involving SGM if science demonstrates a biological link for gender identity (only the non-heterosexual association survives multiple testing correction at FDR < 0.05).

Finally, the age of the respondent was a significant explanatory factor for attitudes on genetic research involving SGM, with younger respondents trending more pessimistic and older respondents trending more optimistic.

### 3.2 Gender variance and its relationship to tolerance and family history of neuropsychiatric conditions

Our survey included a novel means for respondents to describe their gender identity in a continuous fashion. This three-dimensional gender space (Figure 2a) allowed us to create a continuous measure of gender variance (see methods), which varied both by categorical gender identity (Figure 2b) and categorical sexual orientation (Figure 2c). By calculating this continuous measure of gender variance for cisgender respondents as well as those with non-cisgender identities, we were able to include the entirety of our sample in examining gender variance as it relates to other variables measured (Fig. 2d,e). In doing so, we found that respondents who endorsed either a family history of mental illness or neurodevelopmental disorders (Fig. 2d) showed significantly greater gender variance than those who did not (P < 0.01, Wilcoxan test), in agreement with previous findings [5] [21]. In addition, we also found that increasing tolerance toward SGM and associated behaviors was associated with increasing levels of gender variance of the respondent (Fig. 2e, P < 0.001, linear model).

### 3.3 The impact of science on the views of low-tolerance respondents

Those who most strongly endorse science as a past contributor to their personal increased acceptance of non-heterosexuals are among the most tolerant currently (Fig. 2f). Specifically, among those who “strongly agree” that science has made them more accepting, there is a monotonically increasing trend of tolerance (P<0.001, Fisher’s exact test). Among those who “somewhat agree”, those currently showing moderate tolerance are over-represented (P<0.001, Fisher’s exact test), which may indicate a process of transition from low tolerance to high tolerance.

Those who are currently least tolerant of SGM endorse science as a potential avenue for their own increased acceptance of gender non-conforming individuals (Fig. 2g). These lower-tolerance groups trended toward over-representation in the “strongly agree” group (P<0.1, Fisher’s exact test), and were significantly over-represented in the “somewhat agree” group (P<0.001, Fisher’s exact test).

### 3.4 Direct statements from stakeholders

To supplement our objective, quantitative analysis of survey respondent attitudes, we included representative excerpts of stakeholder feedback from the open-text fields at the end of our survey. These statements, included in Table 4, provide important insight into the necessary considerations when conducting genetic research involving SGM.

## 4 Discussion

This study provides the first systematic look at community attitudes toward genetic research at the intersection of sexual orientation, gender identity, and mental health. Past genetic research on sexual and gender minorities (SGM) has often proceeded in the absence of input from stakeholders, and as the field accelerates, it is vital to devote time and effort to engage stakeholders as partners rather than subjects. Our findings suggest that the key predictors of attitudes toward genetic research specifically on SGM are 1) general attitudes toward genetic and mental health research 2) tolerance of SGM and associated behaviors 3) non-cisgender identity and 4) age of the respondent. Importantly, non-cisgender stakeholder status showed a detectable, but ultimately after FDR correction not statistically significant association with pessimism toward genetic research on SGM. Despite these concerns, our findings provide evidence suggesting that stakeholders are willing to engage with genetic researchers and that trust may be earned through that engagement.

A central point of discussion is the increased prevalence of neuropsychiatric conditions within the LGBTQ+ community and how these stakeholders feel regarding genetic research at this intersection. Our data showed that regardless of stakeholder status, the most prominent predictors of attitudes toward SGM genetic research specifically are their general attitudes towards genetic research and mental health research. Non-cisgender identity was to a lesser extent a predictor of these attitudes, but non-heterosexual identity did not achieve significance. Because there is evidence that stakeholder status influences how this line of research is viewed, it is important to emphasize clearly that non-heterosexuality and gender variance are not neuropsychiatric conditions, despite having been pathologized in the past. Homosexuality was removed from the Diagnostics and Statistics Manual (DSM) in 1973 [8] and Gender Identity Disorder [3] was removed in 2013 with the publication of DSM-V [4]. This most recent DSM does include gender dysphoria, which is the diagnosis commonly required in order for transgender individuals to have gender-affirming medical care. Our data showed an association between reported family history of a mental illness or a neurodevelopmental disorder and higher gender variance (this was including both cisgender and non-cisgender respondents). A possible genetic interpretation of this observation is pleiotropy. However, this finding does not necessarily endorse a genetic relationship between the two, and a likely confounding variable is the degree of openness by the respondent when asked these sensitive questions about familial neuropsychiatric conditions and divergences from traditional gender norms.

Although stakeholders are the primary focal point of this study, it was vital to include cisgender and heterosexual respondents (i.e., non-stakeholders), because many of the fears and concerns on the part of stakeholders have to do with how the findings of genetic and other scientific research are received by the broader public. Encouragingly, we found that scientific advancement was reported as a potential pathway toward greater personal acceptance by respondents who also reported the lowest levels of current tolerance of SGM and associated behaviors. In other words, although some stakeholders reported fear of greater stigma and persecution in the face of genetic research on SGM, those would-be persecutors reported that they would be more understanding of SGM if science provided a biological basis for behaviors and identities they don’t currently understand.

### 4.1 Language regarding sex

The use of the term “recorded sex” instead of “natal sex” or “assigned sex” primarily arose from survey feedback and interactions with our community advisory council. These interactions suggested valid objections to both “natal sex” and “assigned sex”. Recorded sex, with its emphasis on the generation of a vital record, i.e., the birth certificate, is our attempt at harmonizing past genetic and other biomedical research with the inclusive, sensitive language that is appropriate for a modern and complex understanding of sex. Although assigned sex is becoming more widely adopted in clinical practice, with the DSM-5 using the term in their language regarding gender dysphoria [4], further research and consideration are needed to develop language that is appropriate in a biological research context and that is not problematic from the perspective of any gender identity.

### 4.2 Limitations of this study

This study is a first important step in the engagement of the SGM stakeholder community in genetic research. Consequently, it is important to consider the limitations of our sample and design, so that results are not over-interpreted. First, our sample skews young, white, recorded female, and highly educated (when compared to national demographics). If this study were replicated in other, more ethnically diverse locations in the U.S., it is possible that some conclusions would be influenced. There are sub-threshold trends in our data that suggest that racial and ethnic minorities are more skeptical of research in general than the white majority of our sample. Despite this limitation, our sample is likely representative of the “samples of convenience” that are often the norm in current genetic research.

### 4.3 Recommendations for genetic researchers

After synthesizing the survey results, the open-text feedback, and interactions that have resulted through re-contact of survey respondents who volunteered for follow-up communication, a number of recommendations have emerged for scientists interested in pursuing research in this area. A common concern was the connection between the eugenics movement, medical research, and SGM. The eugenics movement rationalized abhorrent practices such as forced sterilization, psychiatric institutionalization, and immigration restriction based on traits or identities deemed undesirable by the movement [20], including the LGBTQ+ community. Given the often intersecting history between psychiatry and the eugenics movement, it is incumbent on researchers to plan, execute, and disseminate research in a way that ensures that the basic human rights of SGM are preserved and history is not repeated. The following recommendations are given in the hope of helping researchers achieve this higher standard of more responsible and considerate research.

First, we recommend that all genetic research projects involving SGM have a community advisory council (CAC) composed of stakeholders representing a variety of gender identities and sexual orientations. A CAC can provide input at the study design phase, giving important insight into research questions that are meaningful from a stakeholder perspective. CAC members can also give feedback during the manuscript preparation phase, so that a proper balance is struck between scientific accuracy and considerate messaging. CAC members can also act as community liaisons during the process of fielding community feedback after a manuscript is published.

Second, we recommend that a section of the lab or consortium website be produced, written for a lay audience, that gets out in front of sensitive issues and that clearly answers questions regarding the research motivation and expected results. This website should link to peer-reviewed literature that provides supporting information. The website should also provide contact information for those wishing to express their feedback. Two example websites can be found at http://gender.devgenes.org and https://geneticsexbehavior.info.

Third, publication of results should follow careful preparation of messaging, ideally in the form of a press kit prepared in collaboration with journalists or public relations professionals. One of the main findings of the current study was clear concern that the public would easily misinterpret results of genetic research on SGM. It is therefore incumbent on researchers in this area to not only prepare manuscripts for a scientific audience, but also clear and concise takeaways for the broader public. This should not be left to chance.

Finally, diversity within a scientific team is a tremendous asset. Although there is nothing that would prevent cisgender, heterosexual scientists from performing rigorous and sensitive research in this area, the inclusion of team members who have a personal investment can provide invaluable perspective throughout the research process. It is not clear whether or how this diversity should be signaled, both within scientific circles and to the general public. Such signaling might provide increased credibility, but it also runs the risk of tokenizing individuals. Representation of stakeholder groups among the scientific team should, however, not be misinterpreted as a “blank check” to pursue research in this area without seeking broader, more systematic stakeholder input. Our findings speak to the heterogeneity of perspectives within these groups, and stakeholder scientists are unlikely to adequately capture all the relevant considerations by themselves.

## 5 Conclusion

Our survey sample, highly enriched for LGBTQ+ community members, revealed that attitudes toward genetic research involving SGM are driven by a variety of factors, primarily by general attitudes toward research broadly, as well as tolerance of SGM. To a lesser extent, stakeholder identities were also associated with attitudes toward this area of research, both in optimistic and pessimistic ways. It is important to note that these attitudes were heterogeneous across and within different stakeholder groups. Our data do not support monolithic, cohesive, homogeneous attitudes among stakeholders, although that was a working hypothesis that we considered. Rather, our findings support sexual orientation and gender identity as important but not uniquely decisive factors in how individuals feel about research on one aspect of their identity. We hope these results are understood not as an unequivocal endorsement of this line of research, but instead as a call for engagement and partnership between experts and stakeholders in navigating this challenging frontier.

## 6 Acknowledgements

This version of this manuscript has been read and deemed acceptable by our stakeholder community advisory council, and we thank them for the time they invested in this process. We would like to thank those who gave feedback during development of the survey and manuscript, including Hana Zaydens, Katherine Imborek, Cassie Omlstead, and other contributors who wish to remain anonymous. We would also like to thank Lea Davis and Jess Ehrenfeld for assistance with recruitment. We also wish to thank the survey respondents for making this research possible.

